# Chronic nicotine exposure alters sperm small RNA content in a C57BL/6J mouse model: Implications for multigenerational inheritance

**DOI:** 10.1101/2022.04.27.489636

**Authors:** Dana Zeid, Thomas J. Gould

## Abstract

Multigenerational inheritance is a non-genomic form of heritability characterized by altered phenotypes in the first generation born from the exposed parent. Multigenerational factors may account for inconsistencies and gaps in heritable nicotine addiction vulnerability. Our lab previously found that F1 offspring of male C57BL/6J mice chronically exposed to nicotine exhibited altered hippocampus functioning and related learning, nicotine-seeking, nicotine metabolism, and basal stress hormones. In an effort to identify germline mechanisms underlying these multigenerational phenotypes, the current study sequenced small RNA extracted from sperm of males chronically administered nicotine using our previously established exposure model. We identified 16 miRNAs whose expression in sperm was dysregulated by nicotine exposure. A literature review of previous research on these transcripts suggested an enrichment for regulation of psychological stress and learning. mRNAs predicted to be regulated by differentially expressed sperm small RNAs were further analyzed using biological enrichment analysis, which also supported enrichment of gene expression pathways involved in hippocampus-dependent learning. Our findings point to links between nicotine-exposed F0 sperm miRNA and altered F1 phenotypes in this multigenerational inheritance model. Specifically, differentially expressed F0 sperm miRNAs may regulate the previously observed changes in F1 learning and stress. These findings provide a valuable foundation for future functional validation of these hypotheses and characterization of mechanisms underlying male-line multigenerational inheritance.

## 1. INTRODUCTION

In the United States, incidence of recreational nicotine use and associated addictive disorders has risen in recent years [1]. Traits associated with vulnerability to nicotine addiction are highly heritable [2]. However, consequences of *parental* exposure to nicotine beyond those conferred by the inherited genomic sequence and parental care (i.e., multigenerational transmission) are less clear. Evidence from animal models of nicotine-induced multigenerational inheritance suggests broad phenotypic dysregulation in offspring of nicotine-exposed males [3-6]. We previously found that F1 offspring of male C57BL/6J mice chronically exposed to nicotine exhibited altered hippocampus functioning and related learning, nicotine-seeking, nicotine metabolism, and basal stress hormones [7,8]. However, biological mechanisms involved in these multigenerational effects have yet to be fully elucidated [9,10].

Multigenerational inheritance (also called intergenerational inheritance) is one form of non-genomic inheritance characterized by altered phenotypes in the first generation born from the exposed parent. Though this mode of inheritance has long been shown in plants and invertebrates [11], a growing body of literature suggests that it also occurs in mammals and can be triggered by a variety of parental exposures [12]. Recent work in male exposure models points to sperm small RNA as a promising candidate mediator of multigenerational inheritance [13-15]. Altered sperm small RNA has been associated with nicotine and cigarette smoke exposure in studies utilizing targeted expression analyses [16-21]. However, no studies to our knowledge have conducted transcriptome-wide sequencing of sperm small RNAs in a multigenerational nicotine exposure model. At the level of basic biology, these findings contradict fundamental assumptions of traditional inheritance models, calling for an updated model integrating multigenerational inheritance mechanisms. In the context of addiction research, accounting for ancestral exposure may clarify inconsistencies and gaps in heritable vulnerability [22]. For example, vulnerability to the development of nicotine addiction is known to be genetically transmissible, but heritability estimates and behavioral evidence are inconsistent between human cohorts, as well as between animal and human studies [23,24]. Considering the high degree of variability in ancestral drug exposure between human cohorts, many of these discrepancies may be resolved by consideration of multigenerational effects. Mechanistic characterization of multigenerational inheritance is a crucial first step in identifying the precise role of non-genomic inheritance in nicotine addiction. Because parental exposures cannot feasibly be manipulated in human samples, animal models of multigenerational inheritance are of critical importance in the current discovery stages of this research.

Here, we aimed to characterize sperm small RNAs impacted by chronic nicotine exposure in the context of our previous findings in the nicotine-sired F1 generation. Replicating our previous exposure mode, we sequenced sperm small RNA from nicotine-and saline-exposed male C57BL/6J mice. Several miRNAs with sperm expression dysregulated by nicotine exposure were identified. mRNA targets downstream of differentially expressed sperm small RNA were further analyzed for enriched biological pathways. Our findings point to possible links between F0 sperm miRNAs dysregulated by nicotine exposure and altered F1 phenotypes in this multigenerational inheritance model.

## 1. METHODS

### 2.1 Ethics statement

All procedures were conducted at Penn State University. All procedures were performed in accordance with the NIH Guide for the Care and Use of Laboratory Animals and were approved by the Penn State University IACUC committee (protocol number 201800601).

### 2.2 Subjects

Subjects were male C57BL/6J mice, age 8-9 weeks at the time of minipump implantation and age 12-14 weeks at time of sperm isolation. Mice were purchased from The Jackson Laboratory (Bar Harbor, ME) and shipped at approximately 7 weeks of age. Mice were housed in groups of two to four with ad libitum access to food and water. Mice were housed in standard shoebox cages with cob bedding and access to nesting material and a red domed shelter for enrichment. Upon arrival, subjects were allowed five days of undisturbed acclimation to the colony room prior to minipump implantation.

### 2.3 Nicotine exposure

28-day subcutaneous osmotic minipumps (Alzet, Model 1004, Durect, Cupertino, CA, USA) were implanted under aseptic surgical conditions, 3-5% isoflurane anesthesia. Pumps contained either nicotine hydrogen tartrate salt dissolved in 0.9% sterile saline (administered at 12.6 mg/kg/day, freebase weight) or 0.9% sterile saline. Minipumps were removed 28 days after implantation.

### 2.4 Sample pooling and sperm collection

Sperm pellets were pooled during collection so that three subjects from the same treatment group were represented per final sample to ensure sufficient RNA for sequencing [25]. Prior to pooling, the sample size was 24 nicotine-exposed and 21 saline-exposed mice. After pooling, the final sample size was eight nicotine-exposed sperm samples and seven saline-exposed sperm samples. Each pooled sample contained sperm from mice derived from multiple home cages in an effort to counter any effects of clustering.

Sperm was collected on the fifth day after pump removal. This collection timepoint was chosen based on previous findings showing that primary behavioral and physiological nicotine withdrawal symptoms dissipate by this time [26-28]. Cauda epididymides were dissected immediately after CO2 euthanasia. Cauda were briefly rinsed using phosphate buffered saline (PBS), then transferred to a clean dish containing fresh PBS for trimming of fat and excess tissue. Two to three cuts were made into cauda epididymides and tissue was squeezed gently with forceps to liberate motile sperm. The sperm dish was transferred to a dry incubator set to 37°C for 15 minutes. Media containing motile sperm and tissue was then transferred to a fresh tube and placed back in the incubator for a second 15-minute swim up at 37°C. Supernatant containing motile sperm was transferred to a second fresh tube and spun down at 10,000g for five minutes at room temperature. The pellet was then resuspended with fresh PBS to rinse and re-pelleted. The resulting pellet was then resuspended with 500μL cold somatic cell lysis buffer (SCLB; 0.01% SDS, 0.005% Triton-X) to lyse any contaminating somatic cells. Resuspended sperm was allowed to incubate in SCLB on ice for 10 minutes, followed by centrifugation to pellet, resuspension in fresh PBS for a final rinse, and re-pelleting. Supernatant was removed and 200μL RNALater (Sigma-Aldrich, St. Louis, MO, USA) was added directly to the processed pellet, which was then allowed to incubate at 4°C overnight. After overnight incubation, 200μL ice-cold PBS was added to each sample (to counter density of RNALater during centrifugation, as recommended by manufacturer) and samples underwent a final spin down to pellet. Supernatant was removed and the pellet was stored at -80°C. Sperm purification and RNA extraction methods were adapted from a combination of previous studies [25,29-31] in consultation with Drs. Tracy Bale, Oliver Rando, and Mathieu Wimmer.

### 2.5 Sperm RNA extraction

Sperm RNA was isolated using the Qiagen miRNeasy micro kit (Hilden, Germany) with some modifications. Specifically, sample homogenization steps were modified to ensure complete lysis of sperm cells, as follows: 14.3M beta-mercaptoethanol (reducing agent added to promote complete lysis of sperm heads [25]) was added to QIAzol lysis buffer (1:8 ratio). Samples were homogenized in QIAzol + beta-mercaptoethanol using a benchtop rotor-stator set to medium/high for one minute, followed by a one-minute incubation at room temperature to facilitate dissociation of nucleoprotein complexes. Samples were then homogenized again for one minute, briefly vortexed to ensure complete lysis of sperm heads, and allowed to incubate again at room temperature for 5 minutes.

After this point, RNA extraction proceeded per manufacturer instructions. RNA was eluted using molecular grade water and stored at -80°C.

### 2.6 Small RNA sequencing

Library preparation and sequencing were performed by the Penn State Genomics Core. Sperm small RNA was converted to cDNA using the NEBNext Small RNA Library Prep Kit (New England Biolabs; Ipswich, MA) and sequenced as 75nt single-end reads on the Illumina NextSeq 550 platform (San Diego, CA), producing an average of ∼3.5 million reads per sample.

Sequencing read quality was assessed using FastQC ([32], available at http://www.bioinformatics.babraham.ac.uk/projects/fastqc/). Pre-processing, alignment, and differential expression analysis were performed using the sRNAtoolbox suite ([33], available at https://arn.ugr.es/srnatoolbox/). sRNAbench performed adaptor trimming and read alignment to the mouse genome (GRCm38_p5) using default settings. Quality control was set to filter out reads with Phred score <20 using the minimum mean quality score method. The sRNAde tool was used to perform DEseq2 differential expression analysis. Full analysis settings in sRNAtoolbox are detailed in Supplementary file 1. Raw sequencing data files are available at https://doi.org/10.5281/zenodo.6463310.

### 2.7 Enrichment analysis

Enrichment analysis in Ingenuity Pathway Analysis (IPA; v01-20-04, content version #70750971) was performed to accompany small RNA sequencing results. Two analyses were run:

1. Enrichment analysis using mRNA targets of differentially expressed small RNAs. Specifically, the IPA microRNA target filter was used to identify mRNA targets of sperm miRNAs differentially expressed between nicotine-exposed and saline-exposed males. MicroRNA target filter was instructed to query Ingenuity Expert and Expert Assist Findings, miRecords, and TarBase databases for experimentally observed mRNA targets, and TargetScan for high confidence predicted mRNA targets. Moderate- and low-confidence targets were excluded from analyses. For this initial analysis, the microRNA target filter output (1653 mRNAs) was analyzed using the IPA core expression enrichment analysis. Enriched canonical pathways and upstream regulators were interpreted.
2. To identify candidate genes that may play a central role in F1 phenotype expression, mRNA targets of differentially expressed sperm small RNAs identified by IPA’s microRNA target filter were compared to our previously generated differentially expressed hippocampal F1 mRNAs. In the F1 generation bred from NIC-or SAL-exposed males, we previously identified differentially expressed mRNAs in ventral and dorsal hippocampus (VH and DH; [7]). Here, we compared predicted mRNA targets of differentially expressed F0 sperm miRNAs with a list of combined F1 DH and VH DEGs. IPA core expression analysis was performed on the 75 mRNAs overlapping between these two lists. In line with our goal of focusing on candidate targets with a potential role in F1 phenotype expression in this overlap analysis, we restricted interpretation to predicted enriched diseases and functions. Specifically, enriched diseases and functions were probed for terms related to phenotypes we previously observed in F1 nicotine-sired mice: learning/fear conditioning, nicotine preference, nicotine metabolism, stress response/signaling, and hippocampal cholinergic signaling.

## 2. RESULTS

### 3.1 Small RNA sequencing

Small RNA sequencing of sperm RNA from nicotine- and saline-exposed mice revealed 16 small RNAs differentially expressed (<.05 Benjamini and Hochberg adjusted significance) between the treatment groups (Table 1; a full list of differentially expressed small RNAs is available in Supplementary File 2). All transcripts found to be differentially expressed were annotated as microRNAs (miRNAs). Below, we discuss the four miRNAs differentially expressed at the highest level of significance (≤.01), miR-122-5p, miR-130b-5p, miR-375-3p, and miR-669c-5p. Highlighted findings related to the remaining differentially expressed 12 miRNAs are summarized in brief in Table 2. Note that this is a non-comprehensive overview of representative literature on each of these miRNAs. Note also that many early studies did not specify analysis of the 5p- or 3p-arm miRNA variants. Thus, we specify the arm variant only where this information was available.

**Table 1.** Differentially expressed sperm small RNAs between nicotine-and saline-exposed mice.

| Small RNA ID | Log2 Fold Change (relative to SAL group) | Adjusted p-value |
| --- | --- | --- |
| mmu-miR-122-5p | -1.46 | 0.01 |
| mmu-miR-130b-5p | -1.13 | 0.01 |
| mmu-miR-375-3p | -0.58 | 0.01 |
| mmu-miR-669c-5p | -0.8 | 0.01 |
| mmu-miR-128-3p | -0.57 | 0.02 |
| mmu-miR-151-3p | -0.73 | 0.02 |
| mmu-miR-25-3p | -0.48 | 0.03 |
| mmu-miR-141-3p | 0.48 | 0.04 |
| mmu-miR-151-5p | -0.78 | 0.04 |
| mmu-miR-19a-3p | 0.52 | 0.04 |
| mmu-miR-30b-5p | 0.53 | 0.04 |
| mmu-miR-32-5p | 0.94 | 0.04 |
| mmu-miR-465b-3p | -0.39 | 0.04 |
| mmu-miR-465c-3p | -0.39 | 0.04 |
| mmu-miR-672-5p | -0.84 | 0.04 |
| mmu-miR-871-3p | -0.57 | 0.04 |

**Table 2.**
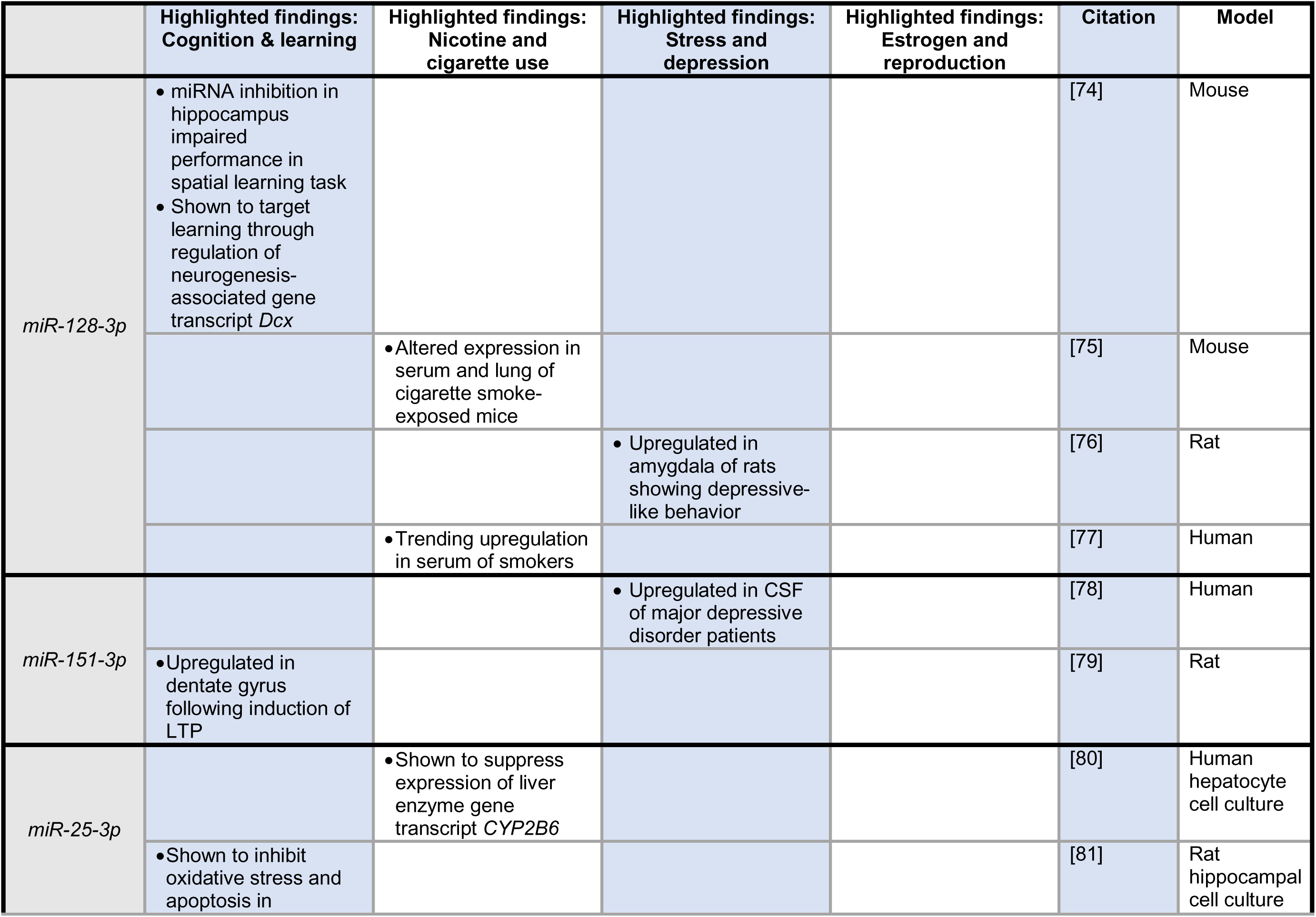
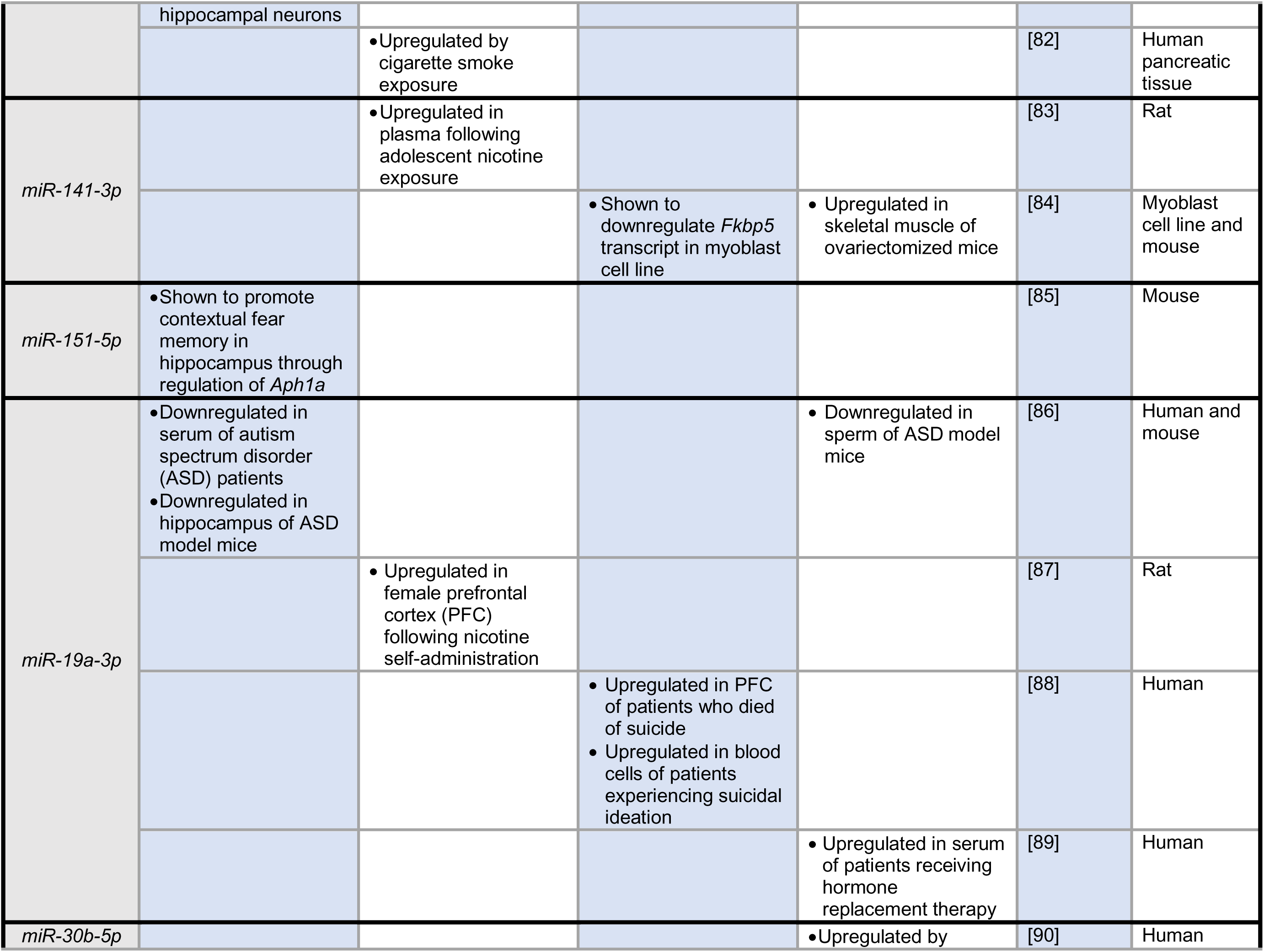
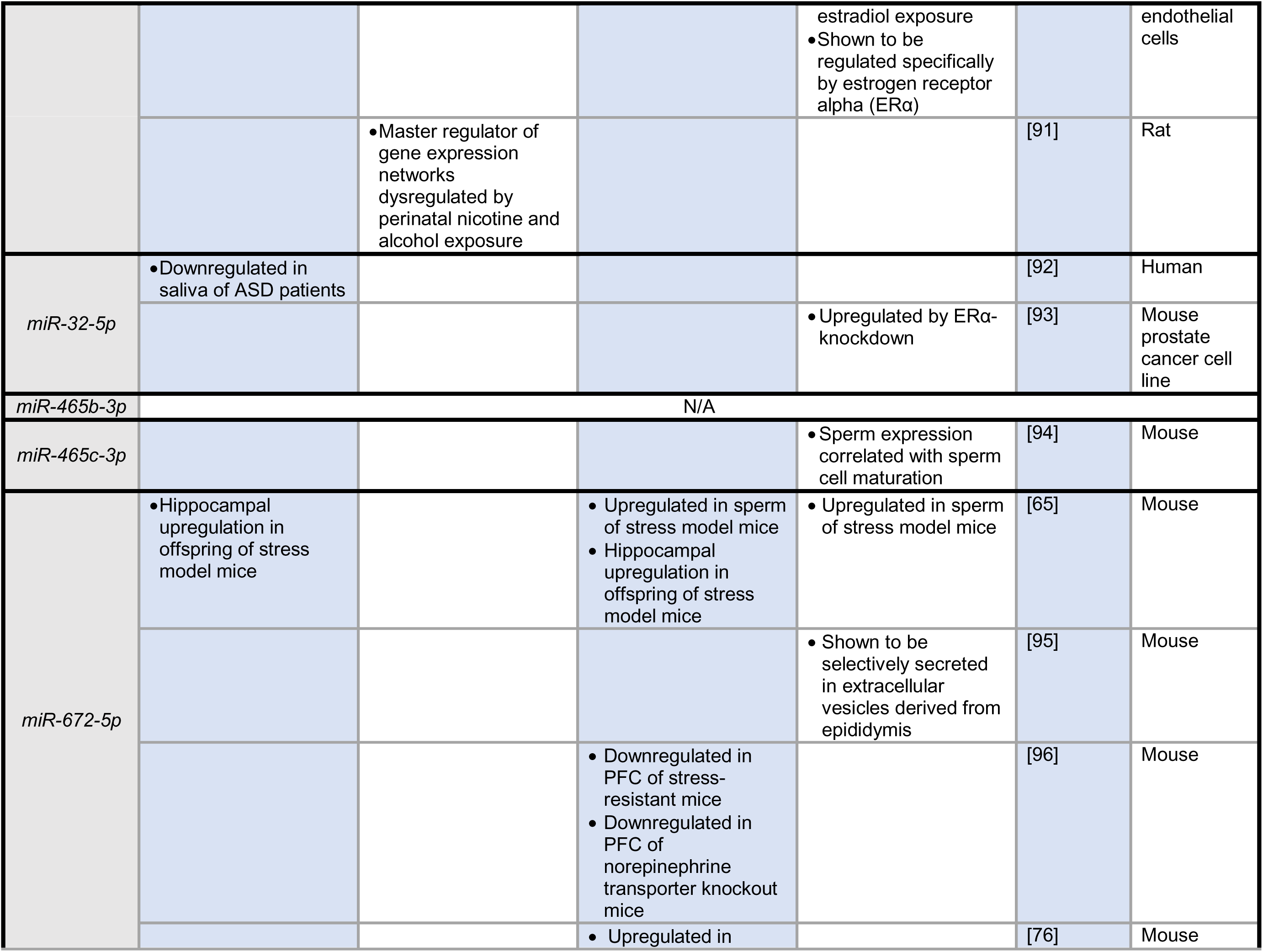
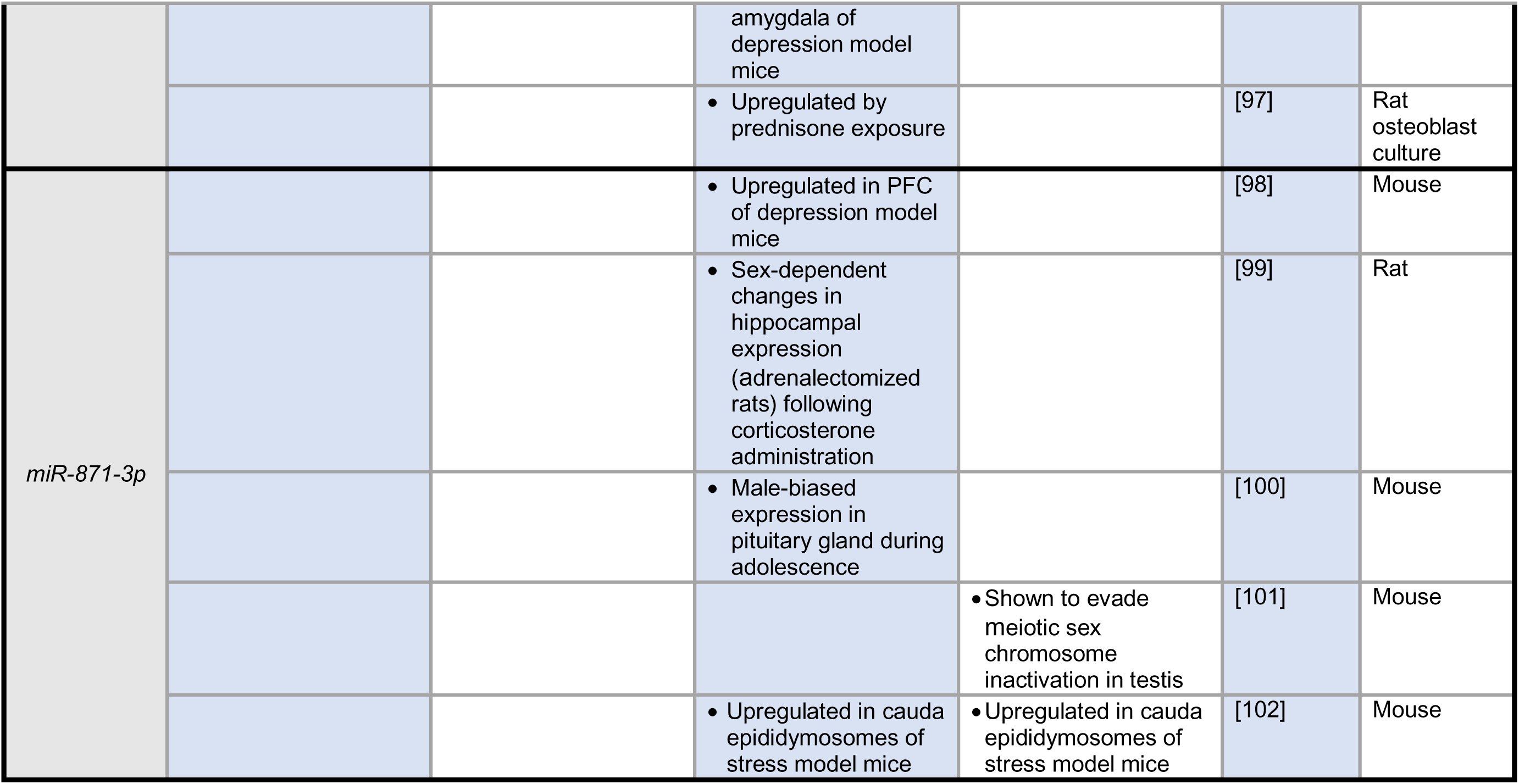
Summarized literature search on last 12 miRNAs differentially expressed between nicotine-and saline-exposed males.

#### 3.1.1 miR-122-5p

miR-122-5p is processed downstream of transcription from miRNA gene *Mir122*, located on murine chromosome 18. MiR-122 was originally identified in mice, with abundant expression in the liver [34]. Subsequent work characterized diverse roles for this miRNA in liver function/dysfunction, including lipid metabolism [35], tumorigenesis [36], circadian cycling [37], and modulation of hepatic viral replication [38]. Later investigations found that miR-122 plays a protective role against ischemic injury in the brain [39]. Relevant to parental exposure models, miR-122 has also been identified as a transcript dysregulated in F1 offspring of female rats fed a high-fat diet [40,41]. One analysis additionally noted increased liver expression of miR-122 in female rats perinatally exposed to nicotine [42].

Interestingly, a recent study identified miR-122-5p as a prominent circulating biomarker of glucocorticoid signaling in two independent human samples [43]. Specifically, Chantzichristos and colleagues initially found decreased levels of circulating miR-122-5p in response to glucocorticoid exposure in a small sample of patients diagnosed with adrenal insufficiency. This association was then confirmed in larger patient populations including individuals with diverse adrenal dysfunction diagnoses and healthy controls exposed to glucocorticoids. Additional analyses revealed a correlation between miR-122-5p and expression of gene *Fkbp5*, a glucocorticoid receptor chaperone.

#### 3.1.2 miR-130b-5p

miR-130b-5p is processed downstream of transcription from miRNA gene *Mir130b*, located on murine chromosome 16. This miRNA has been broadly associated with pathologies in both malignant and protective roles, including multiple sclerosis [44], diabetes [45,46], and several cancers [47-51].

miR-130b-5p has also been implicated in central nervous system functioning. Specifically, this miRNA may be regulated by BDNF (brain-derived neurotrophic factor) and has been identified as a candidate regulator of neural progenitor proliferation during nervous system development [52,53].

#### 3.1.3 miR-375-3p

miR-375-3p is processed downstream of transcription from miRNA gene *Mir375*, located on murine chromosome 1. This miRNA has enjoyed extensive investigation in recent years, revealing association of this transcript with diverse functions and pathologies [54-59], consistent with its characterization as a multifunctional regulator of biological pathways [55]. miR-375 has high expression in the human brain, particularly in the pituitary gland [60], where it regulates adrenocorticotropic hormone secretion [61]. miR-375-3p has also been shown to regulate estrogen receptor signaling [62] and is upregulated by estrogen administration [63].

Relevant to multi- and trans-generational inheritance models, miR-375-3p is concentrated specifically in caput and corpus epididymosomes in the epididymis, suggesting involvement in sperm maturation during epididymal migration [64]. Finally, miR-375-3p was upregulated in sperm of stressed mice and subsequently identified as a candidate causal regulator of transgenerational inheritance in this model [65].

#### 3.1.4 miR-669c-5p

miR-669c was initially implicated in liver aging [66] and immune signaling [67,68]. Later studies suggested a role for this miRNA in neurodegenerative diseases and psychological stress. Specifically, altered brain expression of miR-669c was noted in transgenic mouse models of Huntington’s disease [69] and Alzheimer’s disease [70]. Expression of this miRNA has also been associated with behavioral resilience to chronic stress [71], social isolation stress in a post-stroke mouse model [72], and social defeat fear memory in C57 mice [73]

### 3.2 Exploratory enrichment analyses

#### 3.2.1 Enrichment analysis using all mRNA targets of sperm miRNAs differentially expressed between nicotine- and saline-exposed males

The 1653 mRNAs targeted by sperm miRNAs differentially expressed between nicotine- and saline-exposed males were analyzed using IPA’s core expression analysis. The top five enriched canonical pathways highlighted by IPA were “Pulmonary Fibrosis Idiopathic Signaling,” “Ephrin Receptor Signaling,” “Hepatic Fibrosis Signaling,” “Molecular Mechanisms of Cancer,” and “Wound Healing Signaling.” IPA’s “Upstream Analysis” was used to identify potential upstream regulators interacting with differentially expressed miRNAs. As expected, the top predicted upstream regulators were primarily the differentially expressed sperm miRNAs, as well as miRNAs with similar/identical seed regions. The next top predicted upstream regulators were beta-estradiol and *Esr1*. Supplementary Files 3 and 4 summarize full results of the two IPA analyses, including detailed analysis input parameters.

#### 3.2.2 Enrichment analysis using of differentially expressed sperm miRNA targets overlapping with hippocampal F1 DEGs (differentially expressed genes) between nicotine- and saline-sired mice

Of the 1653 mRNAs predicted to be targeted by differentially expressed F0 sperm miRNAs, 75 overlapped with a combined list of F1 DH and VH DEGs (Supplementary File 2). 11 mRNAs overlapped between all three lists (F1 DH, F1 VH, and F0 sperm miRNA targets). 62 mRNAs overlapped uniquely between F1 VH DEGs and F0 sperm miRNA targets. Two mRNAs overlapped uniquely between F1 DH and F0 sperm miRNA targets. It should be noted that substantial overlap between F1 DEGs and mRNA targets of F0 differentially expressed sperm miRNAs is not necessarily expected, as miRNAs regulate expression of mRNAs acting upstream of complex biological pathways [103]. However, any miRNA targets that do overlap with F1 DEGs may represent hub molecules consistently coordinating multigenerational inheritance from the level of F0 germline exposure to F1 phenotype expression.

Focusing on behavioral annotations in IPA’s diseases and functions analysis, enriched categories included “Cognition,” “Learning,” “Fear Conditioning,” “Conditioning,” “Cued Conditioning,” and “Emotional Behavior.” These enriched behavioral functions were specific to the overlap analysis. This was of particular interest in the context of our previous finding that cued and contextual fear conditioning were enhanced in the F1 generation bred from nicotine-exposed males [7]. That is, it may support the notion that F0 sperm miRNAs inherited by the F1 generation contribute to their altered phenotypes. However, we were concerned that these results simply reflected the nature of our gene enrichment for this analysis – i.e., the 75 mRNA targets of differentially expressed F0 sperm miRNAs were selected based on their overlap with F1 differentially expressed hippocampal genes. This may produce a general enrichment of the hippocampal transcriptome, biasing enrichment terms related to hippocampal functioning. Thus, we reran the IPA core enrichment analysis 10 times using 10 sets of 75 genes randomly selected from the combined list of F1 DH and VH differentially expressed genes, excluding those that overlapped with F0 sperm miRNA targets. The random gene lists were generated using Microsoft Excel’s random value generator (RAND) function.

Our findings using this “permutation” analysis were *suggestive* of non-random enrichment for terms related to fear conditioning in the original analysis (Table 3). Specifically, only in the original analysis were all six of these terms enriched. In five out of ten of the random analyses, none of these terms were significantly enriched. In three of the random analyses, only one of these six terms was enriched. In the remaining random analysis (random analysis #2), three of the six terms were enriched. In all cases but one instance (random analysis #2, “Cued Conditioning”), the number of genes overlapping with the significant enrichment term were greater in the original analysis compared to the random analyses.

**Table 3.** Significant IPA enrichment terms related to learning in original overlap analysis (sperm miRNA targets/F1 hippocampus DEGs; “Original analysis”) and ten randomly selected hippocampus gene set analyses (“Random analyses 1-10”). Significant terms are marked “Yes (#),” with the number in parentheses representing the number of genes in the tested set overlapping with the enrichment term.

|  | Original analysis | Random analysis 1 | Random analysis 2 | Random analysis 3 | Random analysis 4 | Random analysis 5 | Random analysis 6 | Random analysis 7 | Random analysis 8 | Random analysis 9 | Random analysis 10 |
| --- | --- | --- | --- | --- | --- | --- | --- | --- | --- | --- | --- |
| **Cognition** | Yes (10) |  |  |  |  |  |  |  |  |  | Yes (9) |
| **Learning** | Yes (9) |  |  |  |  |  |  |  |  |  |  |
| **Conditioning** | Yes (6) |  | Yes (4) |  | Yes (4) |  |  |  |  |  |  |
| **Cued Conditioning** | Yes (3) | Yes (2) | Yes (3) |  |  |  |  |  |  |  |  |
| **Fear Conditioning** | Yes (2) |  |  |  |  |  |  |  |  |  |  |
| **Emotional Behavior** | Yes (7) |  | Yes (6) |  |  |  |  |  |  |  |  |

## 3. DISCUSSION

Here, we characterized sperm small RNA impacted by chronic nicotine exposure using small RNA sequencing in a mouse model. These findings are of prime significance in the context of drug-induced multigenerational inheritance, which is theorized to be mediated in some capacity by altered sperm RNA content (in male exposure models) [13-15]. Thus, mRNA targets of these differentially expressed sperm miRNAs may illuminate biological pathways mediating phenotypic dysregulation in the F1 generation. Using the same nicotine exposure model employed here, we previously found that offspring of exposed males exhibited altered hippocampal functioning, hippocampus-dependent learning, nicotine-seeking, nicotine metabolism, and basal stress hormones [7,8]. In addition to presenting our comprehensive sperm small RNA sequencing data, we further aimed to interpret these data within the context of our previously published extensive F1 phenotyping.

Using small RNA sequencing of mature sperm cell RNA, we found 16 miRNAs differentially expressed between nicotine- and saline-exposed male mice. The top four differentially expressed miRNAs were discussed, and highlighted findings related to the remaining 12 were summarized (Table 2). Biological enrichment analysis was performed using mRNA targets of these miRNAs to inform hypotheses on mechanisms by which sperm miRNA targets may cluster into functional pathways relevant to F1 phenotype expression.

### 4.1 Enrichment analysis with all mRNA targets of differentially expressed sperm miRNAs

An initial enrichment analysis using all 1653 mRNAs predicted to be targeted by differentially expressed sperm miRNAs suggested dysregulation of pathways related to pulmonary fibrosis, ephrin receptor signaling, hepatic fibrosis, cancer, and wound healing. Of note, our ability to interpret the results of this initial, broad analysis are limited in the absence of soma-wide phenotyping in the nicotine-sired F1 generation. That is, though we and others have characterized several phenotypes dysregulated by paternal nicotine exposure, these were largely restricted to addiction-related behavioral and biological domains. Thus, we present these results primarily for the purpose of hypothesis-generation and comparison with future datasets. A preliminary assessment of the initial enrichment analysis does point to potential congruences with existing literature, however. For example, one group noted increased susceptibility to hepatic fibrosis following injury in F1 offspring of nicotine-exposed male mice [21]. We found altered nicotine metabolism in nicotine-sired mice, and hepatic fibrosis is known to impact nicotine metabolism [104] and be exacerbated by nicotine exposure [105]. Differentially expressed sperm miRNA miR-122’s established role in liver functioning may further link these findings to the current dataset. Relevant to pulmonary functioning, Rehan and colleagues [106,107] noted transgenerational transmission of lung dysfunction and asthma-like phenotypes in a rat nicotine exposure model. Ephrin receptor signaling participates in broad functions throughout the body, but one of its key known roles in the nervous system is regulation of hippocampal learning, which may be in line with our finding of altered fear conditioning in nicotine-sired mice [108,109]. Of potential interest, altered methylation of collagen genes, which participate in wound-healing, was found in sperm of nicotine-exposed rats [110]. Regarding the enriched pathway annotated to cancer processes, the extensive and diverse roles of miRNAs in cancer [111] currently preclude further interpretation of this non-specific enrichment term.

Upstream regulators identified by the enrichment analysis were also assessed, in part to validate target prediction – and, indeed, differentially expressed sperm miRNAs were identified as the top predicted regulators of the target mRNAs. The next top predicted upstream regulators were the endogenous sex hormone beta-estradiol and gene *Esr1*, which encodes estrogen receptor alpha. Though the significance of this predicted regulation by estrogen is unclear within the present model, these findings may be of interest in the context of recent speculation that multigenerational exposures may interact with reproductive hormones to impact the germline epigenome [112].

### 4.2 Enrichment analysis with mRNA targets of differentially expressed sperm miRNAs overlapping with F1 hippocampal DEGs

With the goal of more specifically assessing predicted targets of differentially expressed sperm small RNAs with relevance to our prior phenotyping, we performed an additional enrichment analysis using only predicted mRNA targets that overlapped with our previously generated F1 hippocampal DEGs [7]. For this analysis, we probed predicted enriched diseases and functions, focusing on terms related to phenotypes we observed previously in nicotine-sired mice. Using these criteria, we identified multiple enriched terms related to learning and fear conditioning. To address the possibility that the input gene set (genes expressed in hippocampus) could bias results toward hippocampus-related enrichment terms, we ran ten additional analyses using the same number of randomly selected genes from the overlap list (Table 3). Though we emphasize that the nature of enrichment analysis is exploratory, the random gene set analyses suggested non-random enrichment of learning-related terms in the sperm miRNA predicted targets/F1 hippocampus DEG overlap list. Importantly, this supports the notion that small RNAs dysregulated by nicotine exposure in F0 sperm act upstream of altered F1 learning.

### 4.3 Literature review of differentially expressed sperm miRNAs

A summary literature search of the 16 differentially expressed sperm miRNAs further revealed that many of these transcripts have been linked to functions related to our observed F1 phenotypes. A consistent theme across these findings was the regulation of psychological stress at the level of behavior and underlying physiology. Of note, we previously found that paternal nicotine exposure predicted reduced basal serum corticosterone in F1 offspring [8]. Similarly, Buck and colleagues [3] noted reduced serum corticosterone in female offspring maternally exposed to nicotine. Two of the top four differentially expressed sperm miRNAs, miR-122-5p and miR-375-3p, have been linked to corticotropic signaling [43], and miR-669c-5p was associated with behavioral stress in multiple mouse models [71-73]. Interestingly, one study using a sperm RNA microinjection technique identified miR-375-5p (as well our 15^th^ differentially expressed miRNA, miR-672-5p) as among the transcripts mediating multigenerational inheritance in a paternal stress exposure model. Our 16^th^ differentially expressed sperm miRNA, miR-871-3p, was similarly shown to be upregulated in cauda epididymosomes of stress-exposed males [102]. miR-122-5p, as well as the 8^th^ differentially expressed sperm miRNA, miR-141-3p, were specifically found to regulate *Fkbp5*, a glucocorticoid receptor chaperone. We previously noted altered *Fkbp5* expression and promotor methylation in hippocampus of F1 offspring. As summarized in Table 2, several additional differentially expressed sperm miRNAs were also linked to stress and depressive disorders across diverse models.

Other functions of interest in our literature search were those related to cognition/learning and nicotine metabolism/related signaling. Top differentially expressed miRNAs miR-130b-5p and miR-669c-5p were associated with nervous system development and neurodegeneration, respectively. Among the remaining differentially expressed miRNAs, miR-151-5p was found to specifically regulate contextual fear memories in hippocampus. miR-128-3p, miR-151-3p, and miR-25-3p were identified as regulators of hippocampus-dependent spatial learning/neurogenesis, hippocampal long-term potentiation, and hippocampal apoptosis, respectively. Of potential interest, both miR-19a-3p and miR-32-5p were associated with autism spectrum disorder in human and rodent models.

As mentioned previously, the biological pathway from differentially expressed F0 sperm miRNAs to dysregulated F1 phenotypes is likely multi-tiered, involving several interacting mediators. The regulation of gene expression networks by miRNAs is already known to be profoundly dynamic [103], and this level of complexity is likely multiplied in the context of full mechanistic pathways underlying multigenerational inheritance. Therefore, we hesitate to expound on the directionality of differential sperm miRNA expression in detail. Focusing specifically on studies including sperm miRNA quantification reveals intriguing parallels possibly pointing to stable mechanistic patterns in male-line epigenetic inheritance. We previously noted downregulation of basal stress hormones in the F1 generation bred from nicotine-exposed fathers, and our current analysis revealed downregulation of miR-375-3p, miR-672-5p, and miR-871-3p in sperm of mice chronically exposed to nicotine. These miRNAs were found to be *up*regulated in F0 sperm and cauda epididymosomes in stress-induced multigenerational inheritance models producing heightened stress and depressive phenotypes in rodent offspring [65,102]. Thus, a simple hypothesis supported by these data is that these sperm transcripts act to regulate stress in offspring based on parental experience. Of course, such a theory must be experimentally validated. Further, unlike stress exposure models, the nature of the nicotine exposure in the context of adaptive stress is ambiguous. That is, it is unclear within this theoretical framework whether the presently impacted sperm miRNAs were responsive to nicotine’s direct modulation of stress systems, psychological stress associated with chronic nicotine exposure, or some combination of these factors. Future experiments assessing F0 stress at the time of sperm collection may clarify these dynamics.

### 4.4 Conclusions

We found that chronic nicotine exposure alters sperm miRNA content in a C57BL/6J mouse model. Enrichment analyses and a review of published data on the differentially expressed sperm miRNAs were performed and interpreted in the context of F1 phenotypes previously shown to be dysregulated by paternal nicotine exposure. We conclude that these findings in sum are suggestive of links between nicotine-exposed F0 sperm miRNA and altered F1 phenotypes in this multigenerational inheritance model. Specifically, differentially expressed F0 sperm miRNAs may regulate the observed changes in F1 cognition/hippocampal functioning and psychological stress. These findings provide a valuable foundation for future functional validation of these hypotheses and characterization of mechanisms underlying male-line multigenerational inheritance.

## ACKNOWLEDGMENTS

The authors acknowledge sequencing consultation and services provided by the Penn State Genomics Core. We further wish to acknowledge Drs. Tracy Bale, Oliver Rando, and Mathieu Wimmer for consultation on development of sperm purification and RNA extraction methodology. Finally, we humbly acknowledge the lives of the 45 mice sacrificed to enable this research.

